# Laminar Specific fMRI Reveals Directed Interactions in Distributed Networks During Language Processing

**DOI:** 10.1101/585844

**Authors:** Daniel Sharoh, Tim van Mourik, Lauren J. Bains, Katrien Segaert, Kirsten Weber, Peter Hagoort, David G. Norris

**Affiliations:** Donders Institute for Brain Cognition and Behaviour, Radoud University; School of Psychology, University of Birmingham; Centre for Human Brain Health, University of Birmingham; The Max Planck Institute for Psycholinguistics, Nijmegen; The Erwin L. Hahn Institute, Essen, Germany; Faculty of Science and Technology, University of Twente

## Abstract

Laminar resolution, functional magnetic resonance imaging (lfMRI) is a noninvasive technique with the potential to distinguish top-down and bottom-up signal contributions on the basis of laminar specific interactions between distal regions. Hitherto, lfMRI could not be demonstrated for either whole-brain distributed networks or for complex cognitive tasks. We show that lfMRI can reveal whole-brain directed networks during word reading. We identify distinct, language critical regions based on their association with the top-down signal stream and establish lfMRI for the noninvasive assessment of directed connectivity during task performance.

## Introduction

Top-down and bottom-up information streams are integral to brain function but notoriously difficult to measure noninvasively. In this work, we infer directed interaction between language relevant regions by utilizing functional magnetic resonance imaging with laminar resolution (lfMRI).

Language processing is challenging to understand, as it draws on regions throughout the brain which interact in complex, dynamic configurations. Changes in BOLD amplitude in specific regions have been observed in response to linguistic demands, such as lexical retrieval (left posterior middle temporal gyrus; lpMTG) or the process of combining words into phrases (left inferior frontal gyrus; lIFG (discussed in Hagoort, 2013). During reading, activation in a portion of left occipitotemporal cortex is commonly observed, in addition to activation of language critical regions such as lpMTG (see Carreiras et al., 2014; Price, 2012). Our understanding of such networks would be greatly enhanced if we could elucidate the top-down and bottom-up influences on constituent regions. We show in this work that laminar fMRI can circumvent current methodological limitations which preclude direct, noninvasive measurements of directed interaction.

Anatomists have long been aware of the laminar structure of mammalian neocortex (Campbell, 1905; Brodmann, 1909; von Economo and Koskinas, 1925; Ramon y Cajal, 1911), and have considered its functional implications (see Douglas and Martin, 2004).The advent of retrograde tracers (Rockland and Pandya, 1979) and, more recently, of viral tracing methods has since expanded our understanding of laminar circuits (Barbas, 2015; Rockland, 2017). A key property of laminar circuits is that extrinsic connections to a region tend to target specific layers depending on the hierarchical relationship between the regions. A generalized characterization of this pattern holds that top-down connections preferentially target the deep and superficial layers, whereas bottom-up connections preferentially target the middle layer (Felleman and Van Essen, 1991; Douglas and Martin, 2004, 2007).

With advances in high field MRI, it has become feasible to explore laminar specific functional imaging. A substantial body of work assessing the laminar sensitivity of neurovascular mechanisms indicates that hemodynamic-based measures can be well localized to the site of activation (Duong et al., 2000; Jin and Kim, 2008; Goense and Logothetis, 2006; Goense et al., 2012, reviewed in Norris and Polimeni, in press). This suggests that lfMRI is capable of distinguishing signal driven by neuronal activity in different cortical layers corresponding to hierarchically distinct information streams. Research on this topic has led to reports of task modulated effects at depths associated with the top-down and bottom-up termination sites of mostly sensory (Koopmans et al., 2010; Muckli et al., 2015; De Martino et al., 2015; Kok et al., 2016; Lawrence et al., 2018) and motor cortices (Huber et al., 2017), and at least one laminar resolution experiment has been performed in prefrontal cortex (Finn et al., 2018). In addition, the BOLD signal has been linked to oscillatory activity in different frequency ranges as a function of cortical depth (Scheeringa et al., 2016).

While these previous studies dissociated top-down and bottom-up signal within a region, we demonstrate that these signal streams preserve source information and can be used to identify directed networks.

In this work, we extracted depth dependent time-courses from submillimeter BOLD measurements. We show that reading words compared to pseudo-words increased the top-down BOLD signal observed in the deep layers of the IOTS. The depth dependent responses indicated that different cognitive processes occurred at distinct depths, and would not have been observed at standard resolutions. The depth dependent measurements were then used to identify distinct distributed networks corresponding to top-down and bottom-up signal pathways which targeted the left occipitotemporal sulcus (IOTS) during word reading. The depth dependent signals demonstrated unique connectivity patterns with other regions, thereby establishing the directionality of interaction within the reading network. In addition to the discovery that directed connectivity can be explicitly measured through laminar resolution fMRI, this work provides the first direct evidence that top-down signal to the IOTS is involved in word reading.

## Results

### Top-down task effects in occipitotemporal sulcus

We manipulated the top-down signal directed toward IOTS by visually presenting words and pseudo-words in Dutch (explained in figure 1). Pseudo-words are nonsense words that are orthographically and phonologically legal. As visual representations of words are related to information unavailable from bottom-up sensory information, we expected to observe a relative increase in top-down signal during word reading owing to the linguistic information that could be retrieved from memory (Jackendoff, 2002; Price and Devlin, 2011). The top-down signal was expected to originate mainly in the left temporal cortex (see Hagoort, 2013). Because words are thought to elicit stronger IOTS directed top-down signal, the condition effect, i.e., words against pseudo-words, was expected to target histological layers VI/V, or III/II/I. As discussed, these layers are known to contain extrinsic feed-back targets, and were subsumed within the deep (VI,V) or superficial (III/II/I) bins in our layering scheme (figure 2).

**Figure 1.**
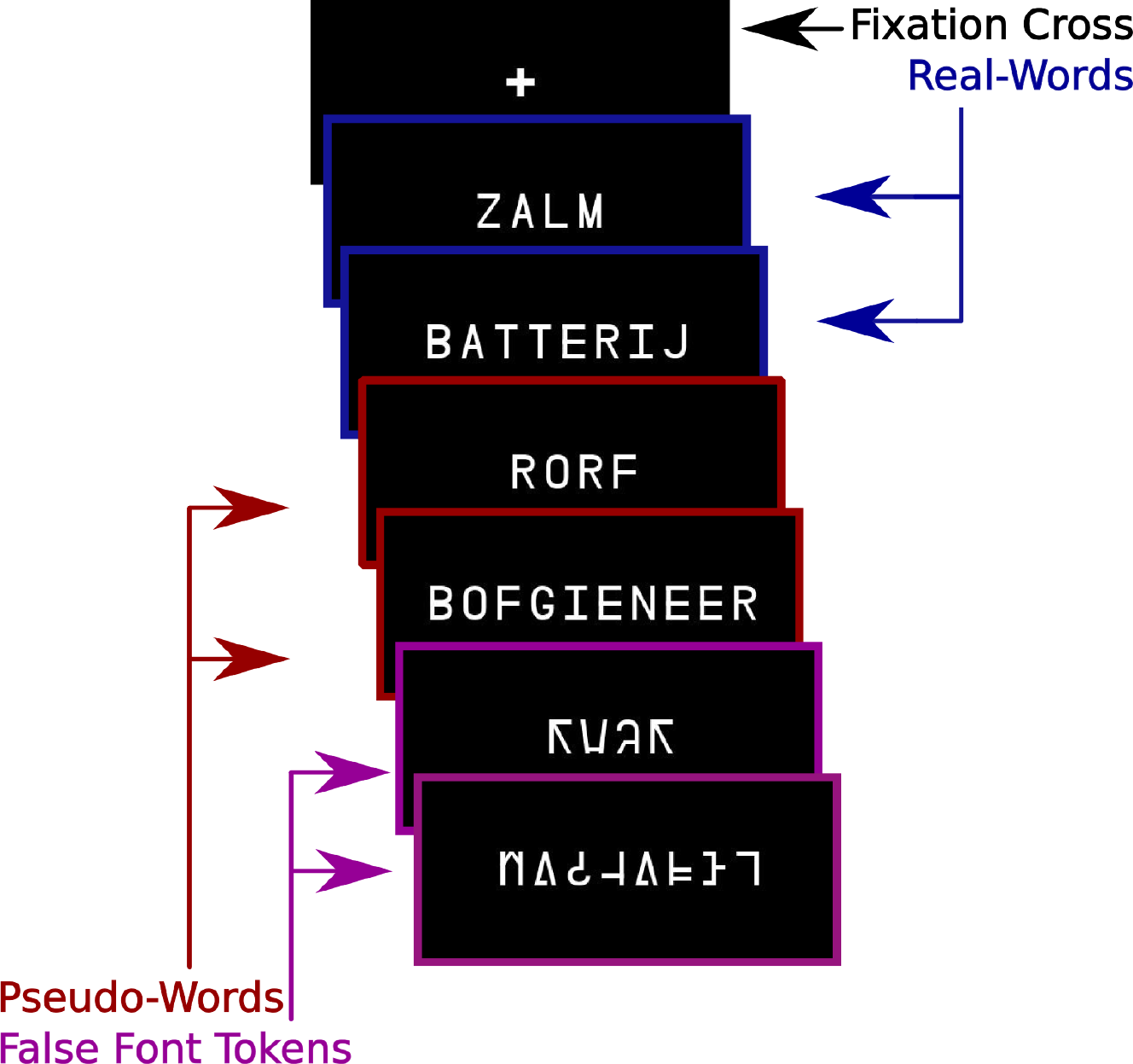
Sample stimuli from each task condition. Word stimuli are in Dutch. English translations of the Dutch word stimuli examples are ‘salmon’ and ‘battery.’ Stimuli were presented in 5 item mini-blocks with the items from the same condition presented in each block.

**Figure 2.**
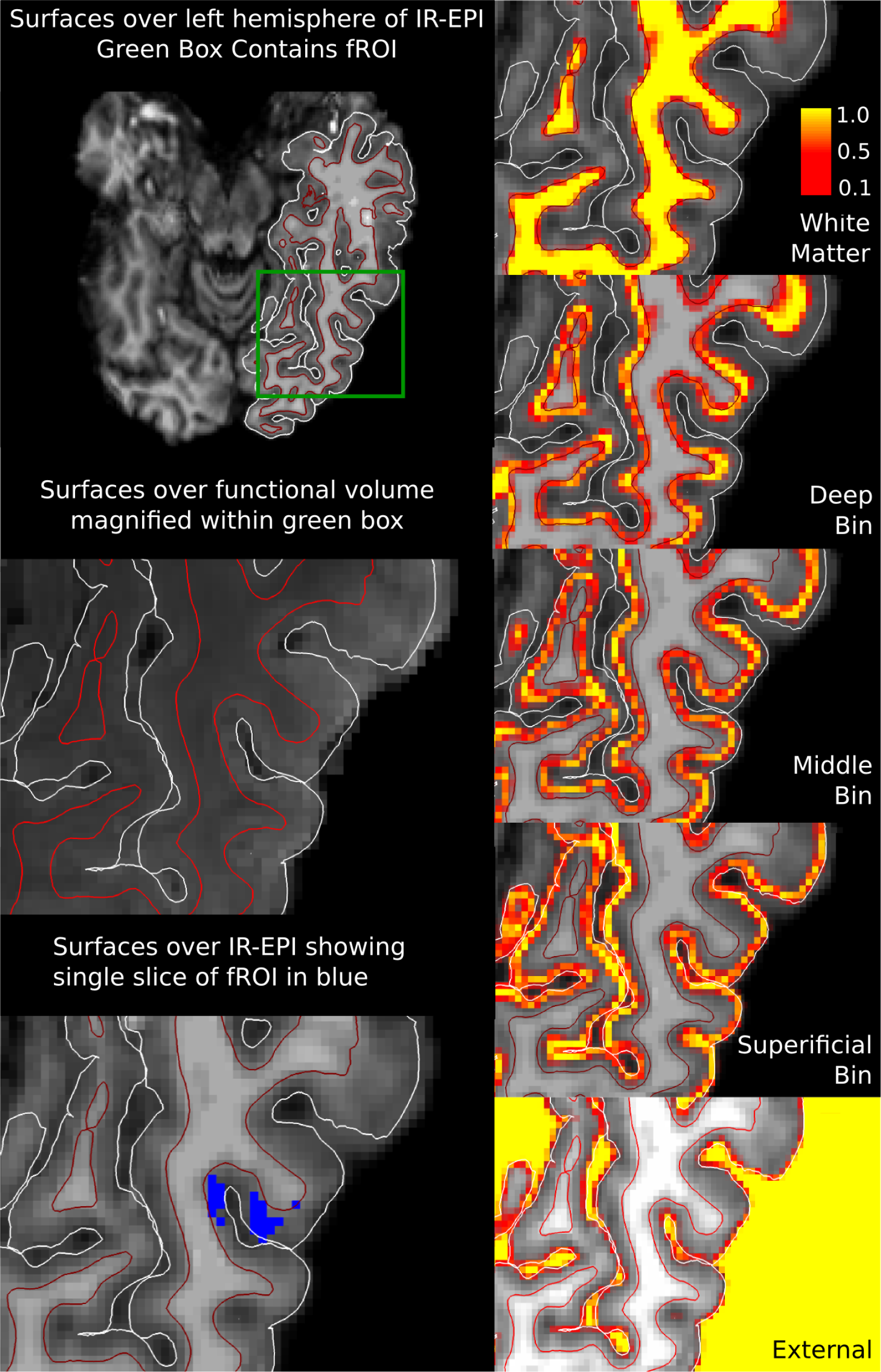
Surfaces and volumetric layering in one subject. Left column: White matter (shown in red) and pial (shown in white) surfaces are shown overlaid on anatomical and T2*-weighted images. Slice through fROI is shown in blue. Magnification within the green box highlights fROI location. Right column: Equivolume depth bins shown in volume space. Each voxel was assigned 5 values between 0-1 according to the proportion of its volume in each of the depth bins. A value of 1 (yellow) indicates that the voxel is fully contained within a given depth bin, 0 (black) that it is not within the depth bin. Intermediate values indicate the fractional volume within a given bin.

Submillimeter gradient echo BOLD 3D-EPI (Poser et al., 2010) data were acquired and analyzed in 22 native Dutch participants during a single word reading paradigm. Participants were instructed to silently read each item presented on the screen. Participants were also presented with items printed in a “false font,” an artificial script designed to conserve the low level features, e.g., angular properties, of the common Latin script. Characters of a false font are letter-like, but visually distinct from characters in orthographies known to participants (figure 1). After some mini-blocks of trials, participants were asked to indicate via button box if the mini-block contained existing words. (Price, 2012).

By contrasting the BOLD response of the false-font items against the words/pseudo-words, it was possible to identify a region that responded preferentially to the orthographic and lexical features of a stimulus rather than to its low-level visual properties. This functionally defined region in the left occipitotemporal sulcus is sometimes referred to as the “visual word form area” (Dehaene et al., 2002). The network properties of this region are of interest, as it is thought that top-down signal directed toward this region may facilitate the process of associating visual information—the sensory information related to the visual representation of the words—with linguistic information (Price and Devlin, 2011). The depth dependent measures reported here were taken from this functionally defined Region Of Interest (fROI).

To analyze the depth dependent BOLD signal, the gray matter volume of the fROI in each subject was divided into three equivolume depth bins following Waehnert et al. (2014), roughly containing the deep, middle and superficial layers, respectively. We restricted our predictions, however, to deeper cortex given that previous work with gradient echo BOLD has discovered top-down effects in the deep bin (Kok et al., 2016; Lawrence et al., 2018). BOLD activity observed in the deep bin was thus taken to best express activity arising from top-down sources, whereas the middle bin was considered to express bottom-up related activity. The depth dependent voxel contributions within the fROI were modeled with a spatial GLM (van Mourik et al., 2019). A temporal, task GLM was then fitted to the depth dependent time-courses that had been computed for each bin using the spatial GLM.

A three-way analysis of variance (ANOVA) was performed to assess the main effects of Depth and Condition (words, pseudo-words) as well as their interaction. Participants were modeled as a random factor. Significant main effects were found for both Depth (*F*(2,105) = 84.37, *p* < 0.001) and Condition (*F*(1,105) = 10.98, *p* = 0.0013). The Depth × Condition interaction was also significant (*F*(2,105) = 12.24, *p* < 0.001). The nature of this interaction was further tested by *t*-statistics on the word and pseudo-word response contrast at each depth bin (figure 3).

**Figure 3.**
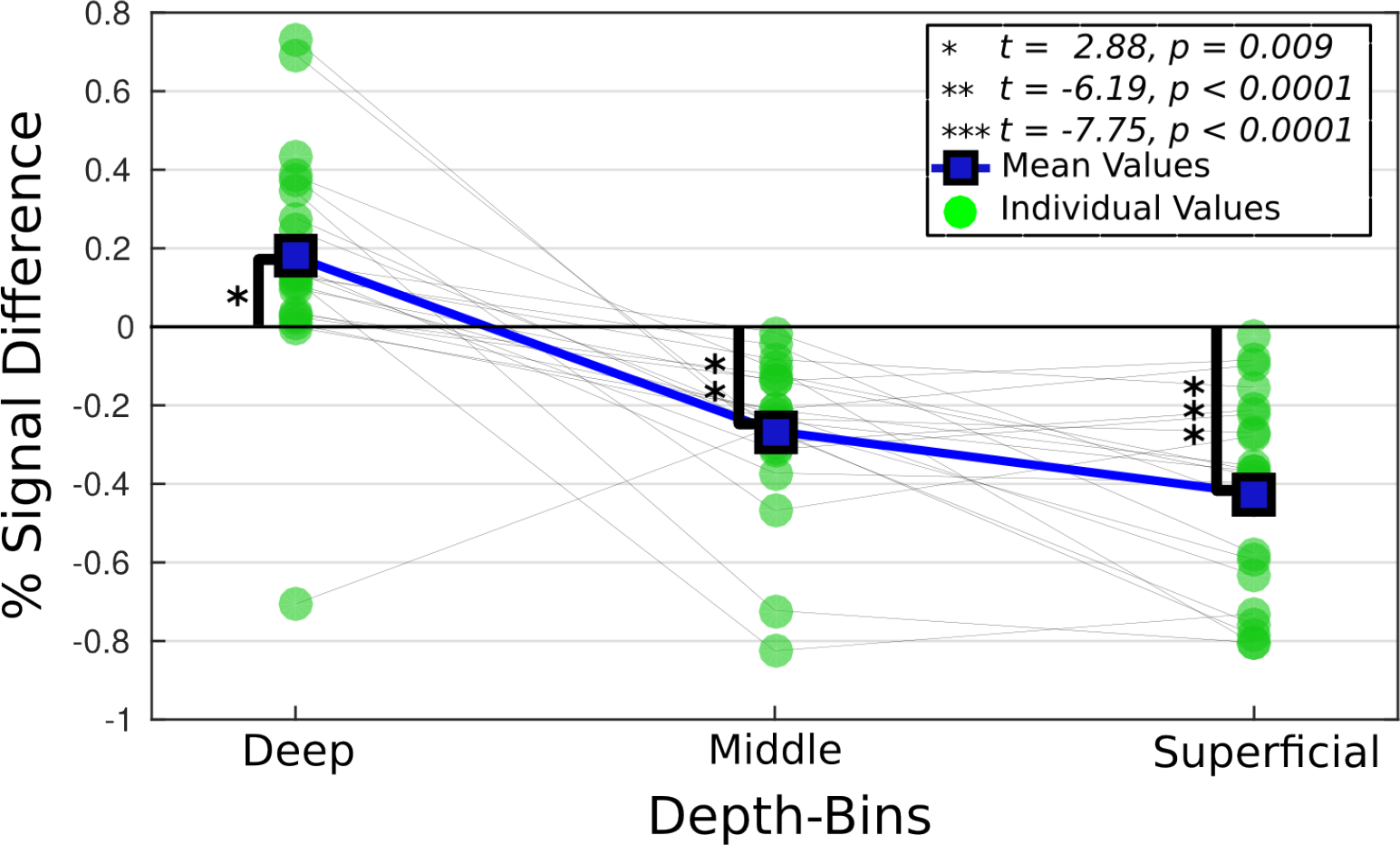
The difference in percent signal change between words and pseudo-words is shown by depth for all participants. Significance bars relate to *t*-statistics computed on the difference from 0 (n = 22)

The depth dependent *t*-statistics were reduced in the middle and superficial bins for words relative to pseudo-words. *T*-statistics in the deep bin were, however, greater for the word condition (figure 3). The greater *t*-statistic in the deep bin is evidence for elevated top-down related signal in the IOTS for word compared to pseudo-word reading. The concurrent reduction of the *t*-statistics within the middle and superficial bins suggests that the increased top-down signal acted to facilitate word reading and was related to the globally attenuated BOLD response throughout the IOTS, as proposed in (Price and Devlin, 2011).

While the relative top-down information content was varied through the word/pseudo-word manipulation, the experiment did not include a bottom-up manipulation. The bottom-up information content was therefore held constant across conditions, and the middle bin effect was unlikely to be the result of differential bottom-up stimulation.

### Within region bin to bin interaction effects during task

Generalized psychophysiological interaction analysis (gPPI) is commonly performed on fMRI data to fit terms in a GLM framework which model the interaction between task conditions (i.e. words and pseudo-words in our experiment) and brain regions (McLaren et al., 2012). The goal is typically to assess the interaction between brain regions in the context of an experimental task. One interpretation of the gPPI analysis is that the strength of the interaction relates to the tendency of the seed region to enhance the response of the target region to a task condition. We used this technique to assess the task-related interactions between the depth bins within the IOTS.

The gPPI model contained all terms used in the first level model with the addition of terms modeling the interaction between the time-courses of the individual depth bins and the task conditions. These were fitted for each participant, and group level *t*-statistics were then computed from the fitted interaction term parameters. The *t*-statistics represent the pairwise effect of each bin on every other bin, and contrasted the word and pseudo-word conditions to determine if inter-bin signal modulation contributed to the lexicality effect shown in figure 3.

Activity in the deep bin was shown to predict decreased activity in the middle bin during word reading compared to pseudo-word reading. This is observed as a negative interaction effect during word reading, indicating that activation in the deep bin predicted a reduced response in the middle bin during word compared to pseudo-word reading, or equivalently, an increased response for pseudo-words compared words. Taken together with the increased deep bin and reduced middle bin *t*-statistics observed during word reading in figure 3, the gPPI results suggest that increased top-down signal during word reading had a suppressive effect on the middle bin, and perhaps on the global signal in the region (table 1).

**Table 1.** Bin to bin intraregional gPPI. Table shows *t*-statistics for contrast of the word compared to pseudo-word conditions. The deep bin shows reduced interaction with the middle bin during word reading.

\* $p \leq 0.01, n = 22$
| Seed and Target | *t*-value | *p*-value |
| --- | --- | --- |
| Deep to Middle | -3.429 | 0.003* |
| Deep to Superficial | -2.194 | 0.04 |
| Middle to Deep | -0.9677 | 0.35 |
| Middle to Superficial | -0.3773 | 0.71 |
| Superficial to Deep | -1.655 | 0.11 |
| Superficial to Middle | -0.8662 | 0.34 |

### Depth dependent whole brain interactions

A second gPPI analysis was performed to examine the effect of the IOTS depth bins on the BOLD response throughout the entire brain. As commonly applied, gPPI analysis is not suited to the study of directed interactions between brain regions. By performing gPPI on depth dependent data, it is, however, possible to leverage knowledge of inter-regional laminar connectivity patterns to obtain directional information. Thus, gPPI on depth dependent data can be regarded as a directed measure, capable of capturing the hierarchical relationship between distinct regions in a distributed network.

The whole brain gPPI model included the interaction terms for the depth bins (deep and middle) and the task conditions (words and pseudo-words). These were then fitted to spatially normalized data in each participant to identify task based, depth dependent networks at the group level. No spatial normalization was applied to the depth bins. The depth bin time-courses (deep and middle) and their interaction (deep × middle) were also modeled and used to assess task-independent connectivity for each depth bin. The interaction between the seeds was included to model intraregional interactions of one bin on another. GPPI targets predicted by the deep bin terms were interpreted as being components of a top-down network related through IOTS. Those predicted by middle bin terms were considered components of a bottom-up network.

Strikingly, the deep and middle bins did not interact with similar brain regions. In the task-independent analysis, we calculated *t*-statistics for the deep and middle bin time-course estimates to obtain a pseudo resting state connectivity measure. Notably, the middle bin time-course experienced strong interactions with bilateral regions posterior to the IOTS, indicative of a feed-forward network including early visual regions. The deep bin interacted with left lateralized gPPI targets anterior to the left OTS, an expected source of top-down afferents to the region. The difference in regional specificity between the two bins is noteworthy. The deep bin interacted with regions within a small volume. By comparison, middle bin interactions were distributed throughout a large bilateral volume (figure 4).

**Figure 4.**
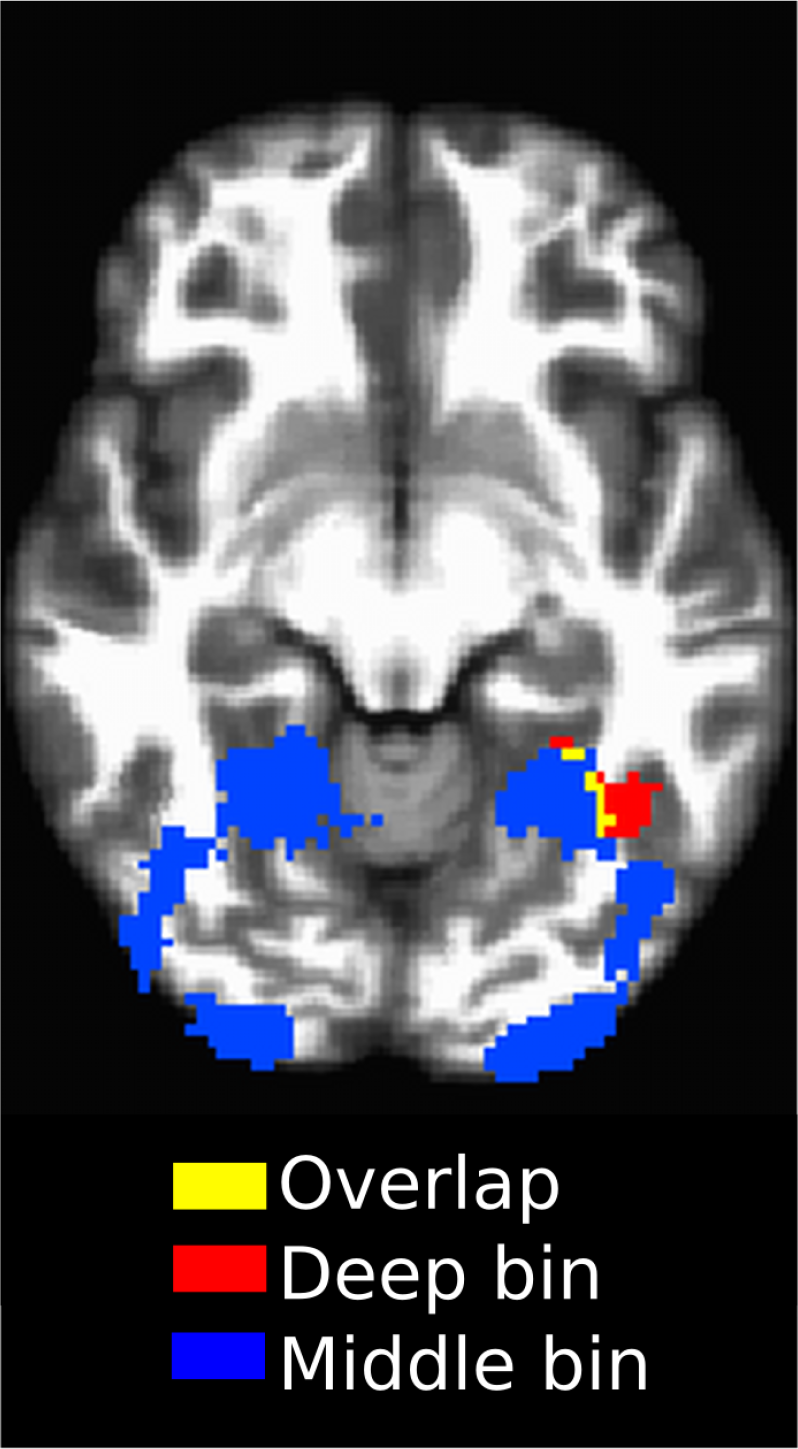
Task-independent connectivity. Colors indicate clusters targeted by the deep bin, middle bin or the overlap. The wide bilateral and posterior targets of the middle bin point to its role as a target of bottom-up signal. The left hemisphere is shown on the right of the image. *p*_*uncorr*_ = 0.001, *α* = 0.05, *n* = 21

The depth dependent interaction terms of the task-dependent analysis identified a top-down network sensitive to the lexicality contrast. The identified network included language critical regions in left temporal cortex and responded preferentially to word reading over pseudo-word reading. When corrected for multiple comparisons (*p*_uncorr_ = 0.001, *α* = 0.05), the deep bin exclusively targeted left middle temporal gyrus (lMTG) and lpMTG, regions commonly associated with lexical retrieval and semantic processing (Hagoort, 2013; Price, 2012; Snijders et al., 2009) (figure 5). The middle bin showed reduced interaction generally, with no clusters surviving correction.

**Figure 5.**
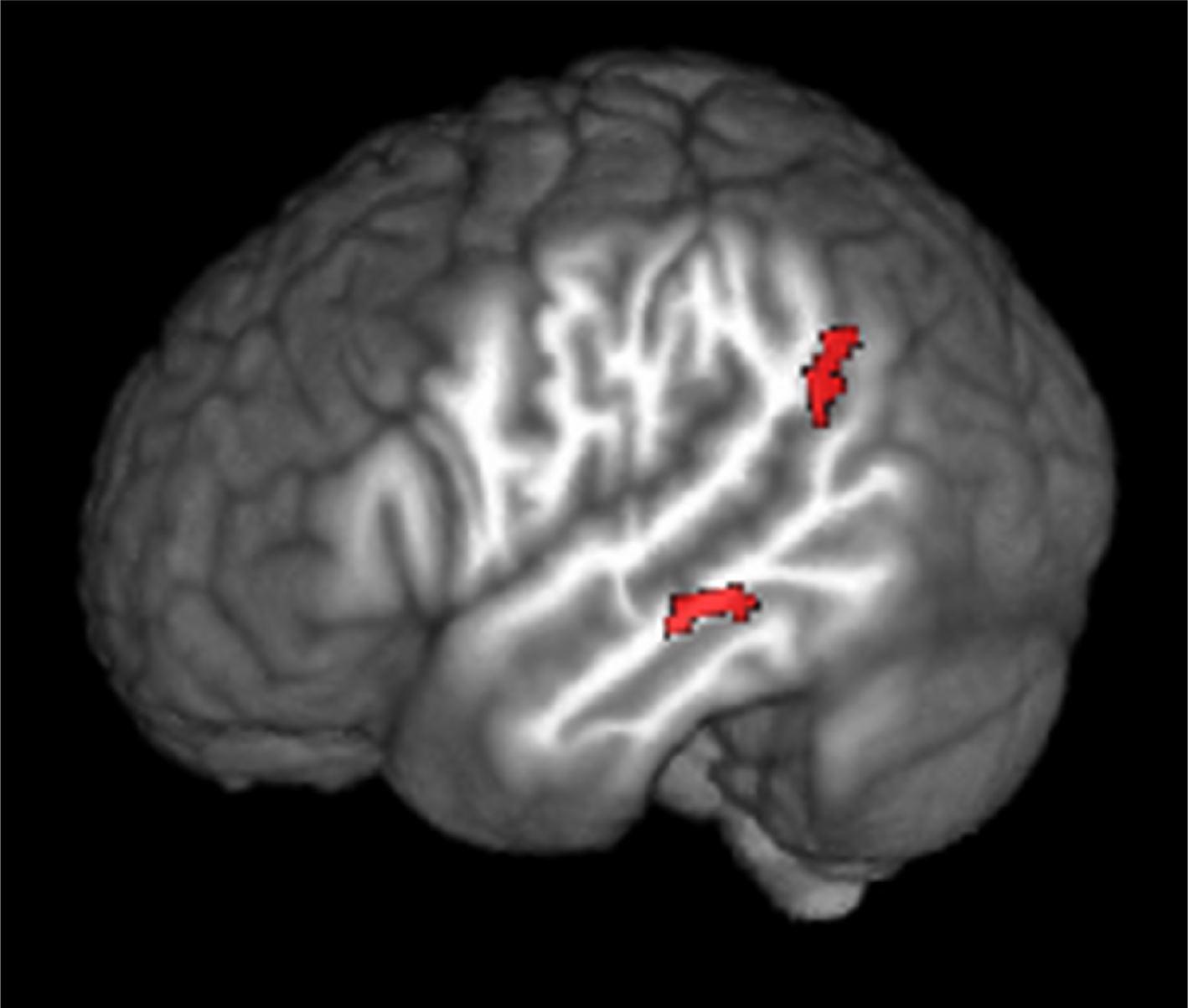
Words against pseudo-words gPPI for the deep bin. Shown in red: the deep bin preferentially targets left lateralized, language critical regions during word reading. *p*_*uncorr*_ = 0.001, *α* = 0.05, *n* = 21

## Discussion

The sensitivity of adjacent depth bins to distinct regions in distributed networks is our most important finding. The unique connectivity profiles associated with each depth bin empirically demonstrate the ability of laminar fMRI to noninvasively identify directed networks. We attribute this outcome to previously unknown characteristics of depth dependent gradient echo (GE) BOLD in combination with gPPI analysis. The findings are direct evidence of top-down interaction between language critical regions in the temporal cortex and the IOTS, relative to the IOTS. Information flow between the temporal and occipital regions is therefore best characterized in terms of top-down rather than bottom-up signaling, a finding which provides insight into the functional role of these regions during word reading. Beyond the unique contribution to the neurobiology of language, the ability to measure directed connectivity has far reaching consequences for expanding our knowledge of brain networks and the processes they implement.

### Depth dependent response to task in IOTS

The reading paradigm used in this work manipulated the top-down information load while holding the sensory input constant across task conditions. The comparatively greater amount of top-down information during word reading resulted in increased top-down signal and elevated BOLD response in the deep layers of the IOTS. The depth dependent *t*-statistics calculated on the contrast of the word and pseudo-word task condition parameters (figure 3) revealed a relative increase in activation in the deep bin during word reading, but a decrease in the middle and superficial bins. The overall response observed when integrating over the three depth bins was reduced, or suppressed, for word reading compared to pseudo-word reading, indicating that the global response did not represent the contrast effect in each bin. This result is evidence that distinct processes occurred within discrete depth compartments during a single condition, and that word reading led to signal increases exclusively in the deep bin. It also illustrates the sensitivity of lfMRI to this nuance which would have been missed by conventional fMRI.

It is not surprising that word reading resulted in a globally suppressed BOLD response in the IOTS given the well-known findings from reading experiments (Price, 2012), but the degree of independence between the deep and other bins is striking. In the presence of global signal reduction, the deep bin expressed increased activation. The relationship between the signal observed in the depth bins to the global signal is interesting and could be uniquely explored in this experiment. This was addressed in the intraregional gPPI analysis and is discussed in the following section.

It is generally agreed that synaptic terminations in layer IV strongly excite their targets and drive spike rate increases throughout the layers of the target region (see Bastos et al., 2012). It was hitherto unknown, however, whether comparatively subtle changes in top-down related responses would be detectable in the BOLD signal in the presence of bottom-up responses. The robust deep bin effect during the word condition indicates that top-down manipulations could indeed be detected without experimentally altering the bottom-up signal, at least with respect to the deep bin. Although it is relatively simple to reduce the bottom-up stream in low-level sensory paradigms, such a restriction would constrain the designs available to researchers investigating higher order cognitive phenomena.

### Bin to bin gPPI

Results from the intraregional gPPI analysis are evidence that the distinct responses within discrete depth compartments arose in part through interactive signal modulation between depth bins.

Observations in the middle bin related not only to the task response, but were also the result of task based interaction with the deep bin. The gPPI analysis revealed a diminished interaction between the deep and middle bins during word compared to pseudo-word reading, which we interpret as suppression of the middle bin driven by the deep bin (table 1). In the depth dependent analysis summarized in figure 3, the deep bin expressed a higher *t*-statistic during word compared to pseudo-word reading, despite the lower *t*-statistics from the middle and superficial bins. We see these results as evidence of top-down connectivity during word reading, but they also suggest that signal in the deep bin acted to suppress signal in the middle bin. When the reduced interaction (table 1) is considered together with the depth dependent *t*-statistics, it is most plausible that the suppression of the global signal during word reading is attributable to the increased deep bin effect.

It should be mentioned that a clear asymmetry is observed when reversing seed/target pairs. The deep and middle bin only interacted such that the deep bin influenced the middle bin, and the middle bin was not observed to influence the deep bin. Given the predictive nature of gPPI there is no reason to expect that reversing the seed/target pairs should return similar results, as would be expected from a correlational measure. As discussed in the seminal PPI paper (Friston et al., 1997), activity in one brain region can be predicted by activity in another on the basis of the contribution from the second region to the first. The PPI method does not imply symmetrical contribution between two interacting brain regions.

In summary, the intraregional results provide evidence in favor of top-down facilitation during word compared to pseudo-word reading (Price and Devlin, 2011), and suggest that activity in the deep layers can act to suppress activation in the middle layers. This mechanistic, inter-laminar description of suppression suggests new possibilities in the use of functional measures to investigate intrinsic connectivity at the mesoarchitectural scale.

### Task independent whole brain connectivity

The connectivity patterns observed in the task-independent gPPI results (figure 4) are in agreement with the expected organization of bottom-up and top-down efferent signal streams in the visual processing hierarchy (Rockland and Pandya, 1979). Relative to a single region, the bottom-up signal is by necessity uninterrupted throughout the processing hierarchy. The top-down stream by comparison does not have these constraints and may be restricted to fewer regions. The larger volume of the middle bin gPPI targets and the smaller volume of the deep bin gPPI targets are consistent with these constraints. The anterior/posterior spatial distribution of the targets is consistent with the top-down/bottom-up signal streams in occipital cortex. Bottom up signal originates in lower regions relative to efferent targets, primarily in visual cortex, for the IOTS. Top-down signal originates in higher regions, often anterior to target populations. Finally, the bilateral middle bin interactions are consistent with the IOTS receiving input from bilateral sources (Chu and Meltzer, 2018).

### Task dependent connectivity

The increased connectivity to the lpMTG and lMTG during word reading was only to the deep bin, which we interpret as evidence of a top-down reading network relative to IOTS (figure 5). No middle bin interactions survived correction. It is remarkable that the different depth bins expressed this degree of specificity and that they could be used to investigate directed interactions instantiating high level cognitive phenomena. The deep bin task interaction is direct evidence that the temporal cortex regions in the reading network relate to the IOTS through top-down rather than bottom-up signal. It is therefore unlikely that the IOTS acts as a node in a forward directed reading network (Dehaene et al., 2002), but rather that word reading is an interactive process whereby top-down signal from temporal cortex to IOTS augments the integration of visual information with linguistic knowledge (Price and Devlin, 2011).

It was not previously known that the commonly used GE-BOLD contrast would be capable of resolving spatially adjacent BOLD responses with sufficient accuracy to interrogate the depth dependent connectivity of distributed networks, or to observe interactions among bins within a region. We attribute these findings to a combination of factors. First among these is the fact that the gPPI analysis regressed out the main effects of the task conditions which best captured variance shared across multiple bins.

Furthermore, previous work has demonstrated that the GE-BOLD response has a peak in the layer in which it originates and a flat tail of far lower intensity up to the pial surface (Markuerkiaga et al., 2016). Variance unique to a given layer will therefore be weaker outside the layer and likely below the threshold for detectability. It follows that some amount of signal contamination will occur from deeper to more superficial layers, but that it should be best represented in the main effects of the task conditions. We argue that the gPPI analysis accounted for downstream signal contamination by regressing out the main effects of the task conditions. The interaction terms could then localize unique variance to the correct depth bins. While it was not the focus of this study, these results further support the notion that the neurovascular response is highly linear and spatially tightly coupled, as previously reported for optical imaging techniques (O’Herron et al., 2016).

This work introduces new possibilities for noninvasively exploring the interaction between brain regions at a spatial scale previously limited to invasive recordings. The benefits of directional connectivity measurements during a task were demonstrated here in the reading network, but are potentially applicable to the study of inter-regional systems throughout the brain. By targeting the reading network, we demonstrated use of laminar fMRI to observe connectivity across multiple non-sensory brain regions during the execution of a uniquely human capacity. It seems likely then that the fundamental approach taken in this paper generalizes to different brain regions and to different topics of inquiry. Distributed networks are known to be critical for brain function, and expanding our study of these systems will ultimately improve our understanding of how higher level functions are instantiated.

## Methods and materials

### Experimental design

#### Task paradigm

Twenty-four native Dutch subjects (13 female, 18-30 years of age, 21 right handed) performed a single word reading task which presented words, pseudo-words and false-font items as task conditions. Subjects 2 and 19 were ultimately excluded from analysis owing to computer failure and signal dropout, leaving 22 data sets for analysis. Data was both lost and corrupted for subject 2 because of computer failure during image reconstruction. It was not possible to recover the lost data. Subject 19 experienced signal dropout in the IOTS and so was excluded from all analysis. Subjects had normal or corrected to normal vision and were screened for reading impairment. Left handed participants were included because regions of interest were determined through functional localization. Language function was observed to be left lateralized in all participants. Informed consent for all subjects was obtained in accordance with procedures of ethical approval of the Donders Centre for Cognitive Neuroimaging and the Erwin L. Hahn Institute.

Initially, the experiment was intended to use a 3 × 2 task design of Lexicality (words, pseudo-words, false-font items) × Length (short, long) with 120 items for each level, and where ‘short’ and ‘long’ designated the length of the words in terms of number of syllables. The length manipulation was intended to modulate the bottom-up load on IOTS. It was determined through piloting that the length manipulation was ineffective and was not analyzed as part of this study. The number of participants was chosen on the basis of previous work using similar acquisition techniques (Kok et al., 2016). Our task thus included three relevant conditions, two of which (words, pseudo-words) were conditions of interest for our analysis, and one of which (false-font items) was used to localize the functional Region Of Interest (fROI) for analysis.

#### Item creation

Word items were selected from a list of high frequency, concrete Dutch nouns taken from the Celex database (Baayen et al., 1995). Words were selected to maximize frequency, minimize the standard deviation of word frequency, and minimize standard deviation of these values for short and long items.

Pseudo-words were generated using Wuggy (Keuleers and Brysbaert, 2010). Pseudo-word generation was constrained on the basis of phonemic neighborhood density, consonant/vowel structure and the number of characters of the word items. The word and pseudo-word stimuli, and the parameters used in Wuggy to generate the pseudo-word stimuli can be found in supplementary materials (StimulisList.xlsx).

False-font items were created by rendering the word items in the false font. The false font (Cohen et al., 2002) was designed to preserve the low level features of familiar orthographically legal characters, but to be visually different from letter shapes. These items are included in supplementary materials (FalseFontItems.pdf). Collapsing across the length manipulation, there were 240 items of each stimulus type. Sample items can be seen in figure 1.

#### Stimulus presentation

Items were presented during fMRI measurements taken over 12 runs. Runs were delimited by breaks in data acquisition. Twenty items of each stimulus type were presented per run, with 60 items presented in total per run.

Individual stimuli were visually presented for 800ms in the center of the display. One item was presented per trial. Items were rendered in white on a black background, as shown in figure 1. Presentation onset was jittered about the 3.9 second TR based on the design optimization calculations obtained using optseq (Dale, 1999).

Stimuli were presented in 5 item mini-blocks in which all 5 items were of the same condition type. Each mini-block was followed by a fixation cross presented for the duration of one trial. On three pseudo-random occasions per run, a question mark was presented that instructed the participant to indicate via button-box whether the previous mini-block contained existing Dutch words. Button-box responses were not analyzed and were considered only to ensure participant compliance. Prior to the experiment, subjects were briefed on the type of items they were to see and instructed to silently read the items on the screen.

The experiment was performed using Presentation ® software (Version 16.1, Neurobehavioral Systems, Inc., Berkeley, CA, www.neurobs.com). Two versions of the experiment were created with roughly half of the participants assigned to each version. The versions differed in block and item order. The different experiment versions were intended to capture latent, unintended effects inherent in presentation order or other version specific properties.

#### Task design model

The task design matrix included condition regressors, temporal and spatial dispersion derivatives, physiologic regressors, motion regressors produced by using SPM version 12 (Penny et al., 2011), drift terms, frequency filters, outlier censors, and constant terms modeling the mean signal per run. Outlier time points were determined using *3dToutcount* in AFNI (Cox, 1996). We considered a voxel to be an outlier if the probability of the distance of its intensity value from the trend exceeded *p* = 0.001 as defined by its location within a Gaussian probability distribution. Time points were excluded from analysis if 2% or more of the voxels at that time point were categorized as outliers.

Stimulus onsets were modeled as instantaneous events with zero duration and convolved with the canonical hemodynamic response function. Condition regressors were created separately for word, pseudo-word and false-font items; and for the long and short items within each condition. In total six condition types were modeled, but only three were analyzed after collapsing across length.

### Acquisition

#### Functional acquisition

Near whole brain, submillimeter (0.943, 0.900 slice direction) resolution T2*-weighted GE-BOLD data were acquired using a GRAPPA accelerated (acceleration factor 8 × 1) 3D-EPI acquisition protocol (Poser et al., 2010) with CAIPI shift kz = 0, ky = 4 (Breuer et al., 2005; Setsompop et al., 2012); effective TE = 20ms, TR = 44ms, effective TR = 3960ms, BW = 1044Hz/Px, FoV = 215mm × 215mm × 215mm with 112 phase encode steps in the slice direction (100.8mm), *α* = 13°, partial Fourier factor=6/8 in both slice and phase-encoding directions. The first phase encoding gradient was applied in the posterior to anterior direction. An axial slab was collected in each subject and positioned to include the occipitotemporal sulcus. The 10cm slab was sufficient to allow complete brain coverage in several subjects and near complete coverage in the remaining subjects. Data were acquired on a Siemens Magnetom 7 Tesla scanner (Siemens Healthineers, Erlangen, Germany) at The Erwin L. Hahn Institute in Essen, Germany. Informed consent was given by each participant in accordance with institutional guidelines. Functional data consisted per subject of 12 3D-EPI data sets of 77 volumes each, though some sessions were incomplete owing to time constraints or other difficulties. No session contained fewer than 10 functional data sets.

#### Anatomy acquisition

Two anatomic images were acquired in each subject using the MP2RAGE (Marques et al., 2010) acquisition protocol (voxel resolution = 0.75mm × 0.75mm × 0.75mm, TR = 6000ms, TE = 3.06ms, T1_1_ = 800ms, T1_2_ = 2700ms, *α*_1_ = 4°, *α*_2_ = 5°, BW = 240Hz/Px, FoV = 240mm × 240mm with 192 slices (144mm)) and a T1-weighted inversion recovery EPI (IR-EPI) protocol based on the parameters used in the functional acquisition protocol. To create the T1 contrast, the following parameters were modified from the functional acquisition: *α* = 90°, T1 = 800ms, TR = 200ms, TE = 20ms. Example images from each type of anatomic acquisition can be seen in figure 6.

**Figure 6.**
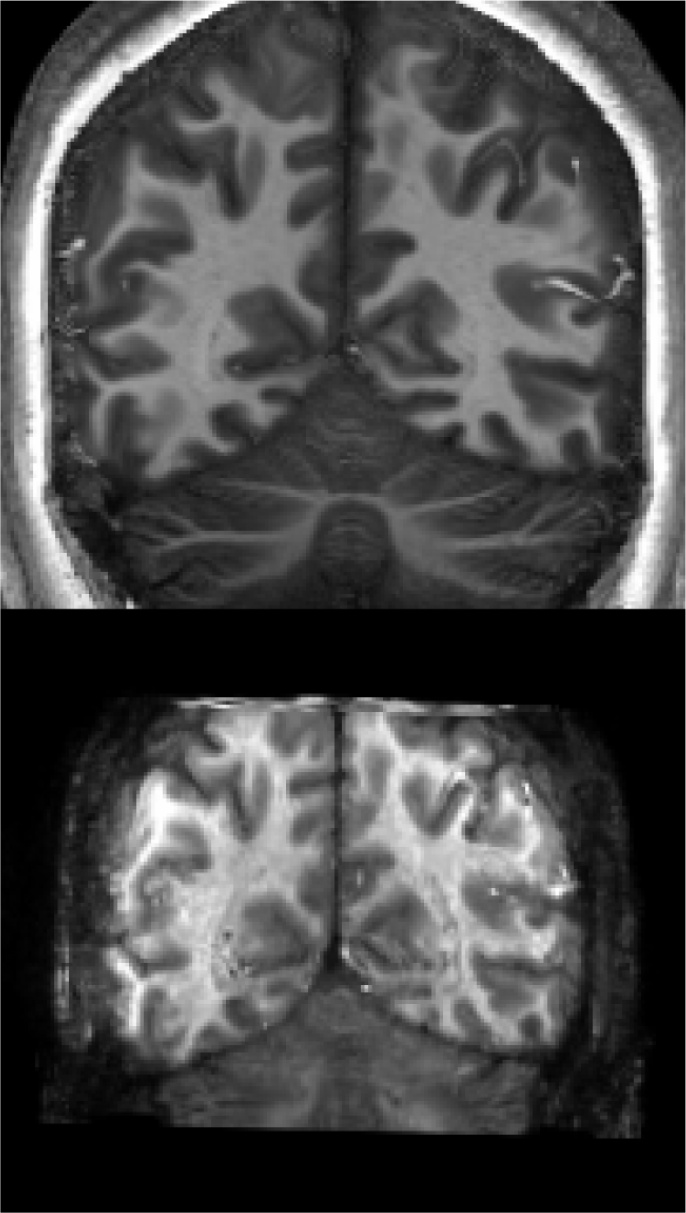
Different anatomy acquisitions used in this experiment. MP2RAGE (top), IR-EPI (bottom). The left hemisphere is shown on the right side of the images.

It was necessary to increase the number of phase encode steps in the slice direction from 112 to 160 and to expand the FoV in the slice direction to ensure fully overlapping coverage with the functional data. The IR-EPI images were used for image registration as they are known to provide high tissue contrast while preserving the geometric distortions of the functional images (Dumoulin et al., 2018). High accuracy, cross-modal registration is challenging, particularly with high resolution acquisitions known to exaggerate geometric distortions. Performing co-registration taking the IR-EPI as the source image mitigated the challenges caused by these distortions.

Field maps were also acquired in some subjects for potential use in distortion correction, though these were not used. The IR-EPI provided sufficient contrast in the native functional space to facilitate high quality registration without the need for distortion correction.

#### Choice of acquisition method for laminar resolution imaging

IfMRI relies on the same neurovascular coupling mechanisms exploited in standard BOLD imaging. In recent years there has been increased support for the concept that the neurovascular coupling occurs at a sufficiently fine scale to make lfMRI feasible (O’Herron et al., 2016). However, the requirement for submillimeter resolution has led to considerable discussion as to the best MR-contrast for interrogating the hemodynamic response.

The standard gradient echo BOLD contrast is highly sensitive to functional activation, but is known to have a considerable contribution from vessels downstream from the site of activation. The less commonly used spin-echo BOLD sequence only acquires data from a subset of the contrast mechanisms that contribute to gradient echo BOLD, but is believed to have a superior intrinsic spatial localization at high static magnetic field strengths (Yacoub et al., 2003; Lee et al., 1999; Duong et al., 2003). In addition, contrasts based on cerebral blood flow and volume (CBF, CBV) should also be considered. It is technically far easier to test and compare these contrasts in animal models, and historically such experiments largely preceded human lfMRI (Goense and Logothetis, 2006; Harel et al., 2006; Kim et al., 2007; Kim and Kim, 2010; Lu et al., 2004; Silva and Koretsky, 2002; Smirnakis et al., 2007; Zappe et al., 2008). The conclusion drawn from these was that CBV was consistently found to have the superior characteristics in terms of spatial resolution, and gradient echo BOLD the poorest. Spinecho BOLD and CBF are somewhere between these two extremes. This hierarchy may be explained in terms of the current view that blood volume changes occur in the arterioles and capillaries (Kim et al., 2007; Behzadi and Liu, 2005; Devor et al., 2007; Hillman et al., 2007), and hence CBV contrast should not be a downstream contrast as is BOLD.

The first laminar fMRI studies in humans are comparatively recent (Koopmans et al., 2010; Ress et al., 2007; Polimeni et al., 2010a), and utilized gradient echo BOLD contrast. Since then, the VASO technique for measuring CBV noninvasively (Lu et al., 2004) has been further developed for application for laminar fMRI at high static magnetic field strengths (Huber et al., 2014, 2015, 2017), and a number of spin echo (Olman et al., 2012; De Martino et al., 2013; Kemper et al., 2016) and combined spin-echo and gradient echo studies (Muckli et al., 2015; Moerel et al., 2018) have been performed. Laminar CBF has to date not been published for human studies. Our rationale for selecting gradient echo BOLD for the current study was based primarily on its exclusive ability to acquire high spatial resolution data from large volumes within an acceptable acquisition time. The two main alternatives – CBV and spin-echo – are currently techniques that are restricted in their volume coverage and suffer from comparatively long acquisition times (Huber et al., 2017; Kemper et al., 2016).

The whole-brain gPPI results we report (figure 5) suggest, however, that GE-BOLD may be capable of more refined spatial localization than previously believed. As discussed in the main text, the ability of the gPPI to account for the task effects likely enhanced our ability to localize signal variance unique to individual depth bins. Simulations from Markuerkiaga et al. (2016) based on reported depth dependent responses in visual cortex to identified a depth dependent peak to tail response ratio of at least 5:1 in all cortical depths at 7T, which would reduce the detectability of unique variance downstream from its source. The gPPI results suggest that this ratio may be conservative, or perhaps influenced by task properties. Our reading experiment presented stimuli at a high frequency relative to presentation rates discussed in Markuerkiaga et al. (2016), which should have produced relatively higher frequency task signal. The vasculature attributed to downstream BOLD effects consists of post capillary vessels draining into larger vessels, whereby differences in the vessel length and flow velocity will act to reduce the coherence of the signal leading the vasculature bed to act as a low-pass filter. High frequency task signal components would therefore be expected to undergo greater attenuation than lower frequency components, and would experience a larger depth dependent peak-to-tail response ratio. In light of our results, it seems clear that unique variance related to each bin was well localized, and that signal contamination was isolated to the main task effects where it could be removed during the gPPI analysis.

### Image registration

High quality registration is critical to laminar fMRI. Given the complications inherent in the registration of submillimeter data, different combinations of tools were necessary to achieve accurate registrations in different participants. The criteria for success were constant, however, across all participants and of an entirely anatomical basis. Alignment quality was determined by visual inspection of brain edges and the left occipitotemporal sulcus. The registration procedure is described in this section.

#### Motion correction

Image realignment was performed using *spm_realign* from SPM 12 (Penny et al., 2011), with parameter values set to achieve the highest quality registration. During this step, a mean functional image was computed to be used as the base image in cross-modal registration.

#### Skull removal

Skull removal was performed on functional and anatomic data prior to cross-modal image registration. Different skull removal procedures were used depending on image modality. The FreeSurfer (Fischl, 2012) watershed function was applied to the IR-EPI data sets, sometimes following a first pass B1 bias field correction (discussed in *B1 Correction*). Nearly all processed brains required manual intervention to remove voxels containing unwanted skull or tissue, or to reintroduce voxels removed in error. Skull removal was performed on all MP2RAGE images in the same manner.

Skull removal was performed on the mean functional images produced during realignment. In this procedure, we manually edited the result of AFNI’s *3dAutomask* program. *3dAutomask* is typically used to remove the skull in images with poor tissue contrast, such as with T2*-weighted images. Parameters for this program were optimized on a per subject basis, and all results were manually edited to ensure that only voxels containing skull were removed. We found these results to be adequate on the basis of visual inspection following manual intervention, where ‘adequate’ describes results which did not contain residual skull or exclude voxels containing brain-matter. *3dAutomask* parameters were iteratively optimized until a result was obtained which reasonably limited the necessary manual intervention.

#### B1 correction

B1 correction on the IR-EPI data was unsuccessful on 5 data sets using the standard tools available in the FreeSurfer suite, resulting in failed skull removal and inaccurate segmentations. We were able reduce problems related to B1 inhomogeneity by applying an additional B1 correction before applying FreeSurfer tools. Our approach was to calculate a firs*t*-pass transform for the mean functional and IR-EPI images (with the skull) and apply the transformation to the mean functional image. We then utilized the B1 bias captured in the mean functional image to correct the IR-EPI anatomic images. Following the initial coregistration, the mean functional image was smoothed and voxel-wise intensity scaled between 0.3*v* and 0.9*v* of its intensity value v to prevent extreme values from unduly influencing bias correction. The IR-EPI was then divided by the scaled image, and the result was taken as the corrected image. The corrected image could then be coregistered to the original mean functional image and used as the input dataset for the standard FreeSurfer processing pipeline. We observed a marked improvement in both the coregistration results and the results of the FreeSurfer segmentation and surface generation after performing this correction (figure 7). Computation time was drastically reduced as well, in some cases up to 15 hours.

**Figure 7.**
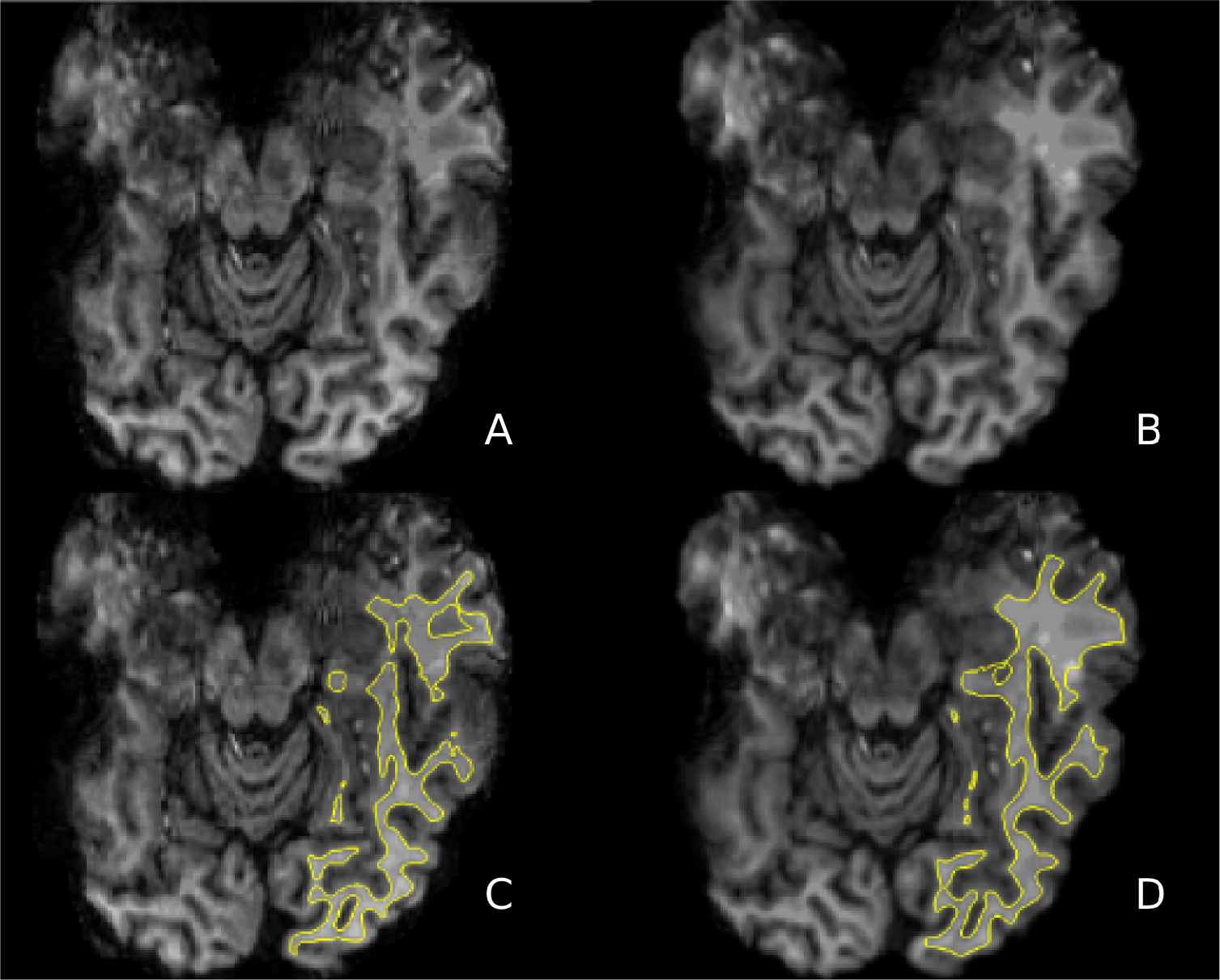
Comparison of FreeSurfer white matter surface generation on the left hemisphere before and after supplemental B1 correction. (A) uncorrected; (B) corrected; (C) surfaces generated from A; (D) surfaces generated from B. The left hemisphere is shown on the right side of the image.

#### Coregistration

Within-subject coregistration was performed using the skull-removed mean functional image and the skull-removed IR-EPI image. Using this image set mitigated registration error owing to image distortion typically observed in EPI acquisitions. High quality coregistration was crucial to the laminar analysis featured in this experiment, as the accurate definition of tissue boundaries in functional space follows only from a highly accurate coregistration of the structural and functional images. Note also that the transformation computed in this step was applied to the structural image to avoid introducing interpolation errors.

Several coregistration tools were used to calculate optimal image alignment. For a given subject, multiple transforms were calculated and visually inspected. The best alignment as determined by visual inspection was taken for further analysis. Volume coregistration was performed using FreeSurfer’s robust, outlier insensitive registration cost function as implemented in *mri_robust_register*. If the resulting transformation resulted in poor registration, we then used the NMI cost function implemented in *mri_robust_register* and finally the NMI cost function implemented in AFNI’s *3dAllineate*. If necessary, manual improvements were applied to the best transformations generated by these tools. Registration quality was assessed by visual inspection of alignment along the left occipitotemporal sulcus and brain edges.

In 11 subjects, failure to reconstruct surfaces from the IR-EPI image made it necessary to perform surface reconstruction on MP2RAGE data, and therefore to bring the MP2RAGE surfaces into register with the functional data.. Following the initial coregistration described above for the IR-EPI images, the IR-EPI images were generally in good alignment with the task data and could serve as the source image for this purpose. In this case, MP2RAGE surfaces were aligned to the IR-EPI volumes using FreeSurfer’s boundary based registration program. If the IR-EPI volume was not in good alignment with the functional data, an initial alignment between the MP2RAGE and functional data was first computed using the tools described in the previous paragraph before performing the boundary based registration. The boundary based registration procedure is described below in a dedicated section.

#### Normalization for group analysis

Functional data for each subject were mapped into MNI space for use in the whole-brain gPPI group analysis. The skull-removed MP2RAGE image was first brought into alignment with the skull-removed mean functional image. After inspecting the quality of the registration, the functionally aligned MP2RAGE images were aligned to the MNI128 template available in standard FreeSurfer installations. This transformation was concatenated with that obtained from the inverted matrix from the initial coregistration and applied to the motion corrected functional data. The result of this spatial normalization was MNI mapped functional data for each subject.

#### Tissue segmentation and surface generation

Tissue segmentation was performed in FreeSurfer using the skull-removed IR-EPI image. Failures to properly reconstruct subject surfaces were addressed by inserting control points, applying additional normalization as described in a previous section, or disabling the correction of defects in surface topology if they did not occur in experiment critical regions. The IR-EPI images commonly included artifacts in noncritical locations that would result in discontiguities in the surface and unsuccessful surface generation. As these defects did not often occur near IOTS, it was possible to generate accurate surfaces even after bypassing correction. If surface reconstruction failed following these interventions, the MP2RAGE dataset was used in place of the IR-EPI, and additional registration steps were applied (discussed below).

#### Boundary based registration

Surfaces reconstructed from the IR-EPI image did not require additional alignment to the functional data beyond resampling the FreeSurfer generated surfaces from “conformed space” to functional space. “Conformed space,” native to FreeSurfer, is a 1mm isotropic 256^3^ grid in the RAS coordinate system.

Surfaces reconstructed from MP2RAGE images underwent an additional registration step using *bbregister*, FreeSurfer’s boundary based registration (BBR) tool. The goal of this procedure was to produce surfaces in register with the functional data. As the IR-EPI and MP2RAGE data sets were generally well aligned from the coregistration procedure described previously, the main purpose of the BBR was to find a solution accommodating the distortions affecting surface placement along the fROI. Using a boundary based cost function, IR-EPI images that were unable to be used for surface generation were aligned with the boundaries generated from the MP2RAGE images. The inverse of this transformation was then applied to the surfaces to align the boundaries to the IR-EPI image. If the alignment generated by the boundary based registration procedure was found to be inaccurate, simple solutions to improve accuracy involved optimizing the registration for the fROI through a weighting mask, manual intervention, or improving the alignment of the two images prior to the boundary based registration. Failing a simple solution, we also computed a nonlinear boundary based registration (van Mourik et al., 2018). In this approach, the registration algorithm recursively divided and aligned surface segments to increase registration accuracy.

The importance of highly accurate image alignment in laminar resolution imaging cannot be over-stated. In the present work, registration inaccuracies in excess of 1mm had the potential to displace entire bins, leading to meaningless results. Great care was taken to ensure accurate registrations and alignment of the surfaces with the functional images. As in the other registration procedures, registration quality was assessed only through visual inspection of key anatomy.

#### Equivolume contouring

The gray matter volume of each subject was partitioned into equivolume bins using the OpenFmri (https://github.com/TimVanMourik/OpenFmriAnalysis) implementation of the equivolume contouring approach described in Waehnert et al. (2014). The equivolume method increases the likelihood that the histological profile of each bin is consistent throughout the given region.

We partitioned the gray matter volumes into 3 bins: the smallest number of bins which allowed for the dissociation of the deep, middle and superficial contributions to the overall BOLD signal. For the purpose of the spatial GLM, it was necessary to include two additional non-cortical bins representing white matter and CSF volumes respectively. The inclusion of these additional bins was due to partial volume effects caused by voxels extending outside the cortical strip. Voxels observed within these boundaries were assigned a value representing the fractional volume observed within a particular set of boundaries.

The ultimate output of this procedure was a 4D dataset whose first 3 dimensions represented spatial coordinates and whose 4th dimension represented the different bins. Incrementing over the 4th dimension indices gave the fractional volume of each voxel found in that particular bin. This volume is referred to as the layer-volume distribution.

In our partition scheme, the volume subsuming the six histological layers was partitioned into three bins. We argue that the histologically coarse bins were sufficient to dissociate top-down and bottom-up signal contributions. At the mesoarchitectural level of lfMRI, it is not practical to measure individual histological layers. One common approach to this challenge has been to consider a simplified model of layer interactions which merges supragranular (layers I,II,III), granular (layer IV) and infragranular (layers V,VI) histological layers into three logical layers based on shared connection tendencies (Koopmans et al., 2010; Muckli et al., 2015; De Martino et al., 2015; Kok et al., 2016; Lawrence et al., 2018). This model is based in large part on patterns of laminar connectivity discovered in Rockland and Pandya (1979) when exploring the link between anatomy and functional hierarchy, and schematized in the Felleman and Van Essen hierarchy 1991. The simplified laminar model has proven valuable when constrained by functional data, and has informed efforts in lfMRI.

### Physiologic noise removal

Cardiac and respiration data were collected concurrently with the functional data using a pulse oximeter and pneumatic belt. Physiologic regressor estimation up to the 6th (cardiac) and 8th (respiration) order was performed using a modified version of the PhysIO toolbox of the TAPAS suite (Kasper et al., 2017). These minor modifications were necessary to account for unique log file formats produced by the equipment at the scan site. Regressors were then included in the design as nuisance regressors. Regressor quality was assessed with a partial *F*-test.

### Non-laminar analysis

The task design was fitted using the generalized least squares regression implemented in the OpenFmri analysis suite. First level *T*-statistics were calculated in MATLAB version R2014B (The Mathworks Inc.) for condition and contrast effects in each subject individually. Several versions of the first level analysis were performed to calculate parameter fits in both native space and MNI space, and with different levels of spatial smoothing applied. This was necessary owing to requirements of the different analyses reported in this work.

Native space data with were spatially smoothed with a 1mm and 3mm Gaussian kernel. Data smoothed with the 3mm kernel were mapped to MNI space and used in the whole-brain gPPI analysis. Data smoothed with the 1mm kernel were analyzed in the first level GLM used to identify the fROIs. Following fROI identification, the fROIs identified in each subject were resampled to native resolution and used as an inclusive mask of the original resolution, native space functional data. The voxels included in this mask then underwent depth-dependent signal extraction.

### Procedure to define functional region of interest

Anatomically, the region of interest was located proximal to the fundus of the occipitotemporal sulcus. It was functionally defined as a cluster of voxels which responded to visually presented words and pseudo-words, but preferentially to pseudo-words. In addition, this region is known to express reduced BOLD amplitude to false-font items compared to items composed of orthographically legal characters (Price, 2012).

The region was defined in each subject through a series of masking operations implemented with AFNI’s voxel-wise dataset calculator *3dcalc*.These operations were performed on the *t*-statistics from the 1mm smoothed, native space analysis. First, voxels were removed if they did not reach threshold in both the word and pseudo-word conditions. This was defined as a *t*-statistic where 1 ≤ *t* ≤ 3. *T* was initially set to *t* = 2 and was increased or reduced if the number of surviving voxels fell outside of the desired range (see below). We then removed all voxels with a larger *t*-statistic for the false-font condition than for either the word or pseudo-word conditions. Finally, voxels were excluded if the difference between *t*-statistics of words and pseudo-words was larger than the original activation threshold. Clusters were considered for inclusion if they were 1) located within the extent of the occipitotemporal sulcus, if 2) cluster size was between 100 and 400 voxels, if 3) 30-50% of the voxels responded preferentially to the word condition over the pseudo-word condition, and if 4) the total response was comparable between the word and pseudo-word preferred voxels when considering the proportion of voxels preferring each condition. These criteria were selected to isolate a functional region which is known to respond preferentially to both words and pseudo-words compared to false-font items, prefer pseudo-words to words, and contain a mixture of individual voxels which prefer each condition. The fROI selection procedure was biased by design toward pseudo-word activation because stronger pseudo-word activation is a functional feature of the region (Price, 2012).

Further considerations were made with respect to the importance of cluster contiguity. Given the high spatial resolution of our data, it was possible to distinguish populations of active voxels spanning the CSF boundary bridging the occipitotemporal sulcus. Following the removal of the voxels located in CSF in some subjects, formerly contiguous clusters became distinguishable. We determined that the most reasonable approach was to include formerly contiguous voxels in the laminar analysis. Given that this region is often functionally defined and generally identified near the fundus of the occipitotemporal sulcus, partial volume effects have almost certainly influenced fMRI measurements at standard resolutions. The decision to exclude populations of voxels stranded on either side of the chasm would have proven arbitrary in that non-laminar studies investigating this region typically lack the resolution to distinguish fusiform and inferior temporal populations. We concluded that allowing for discontiguities in the left OTS fROI more faithfully adhered to the literature definition of the region than an ad hoc justification for voxel removal. Functional ROIs as defined in two participants can be seen in figure 8.

**Figure 8.**
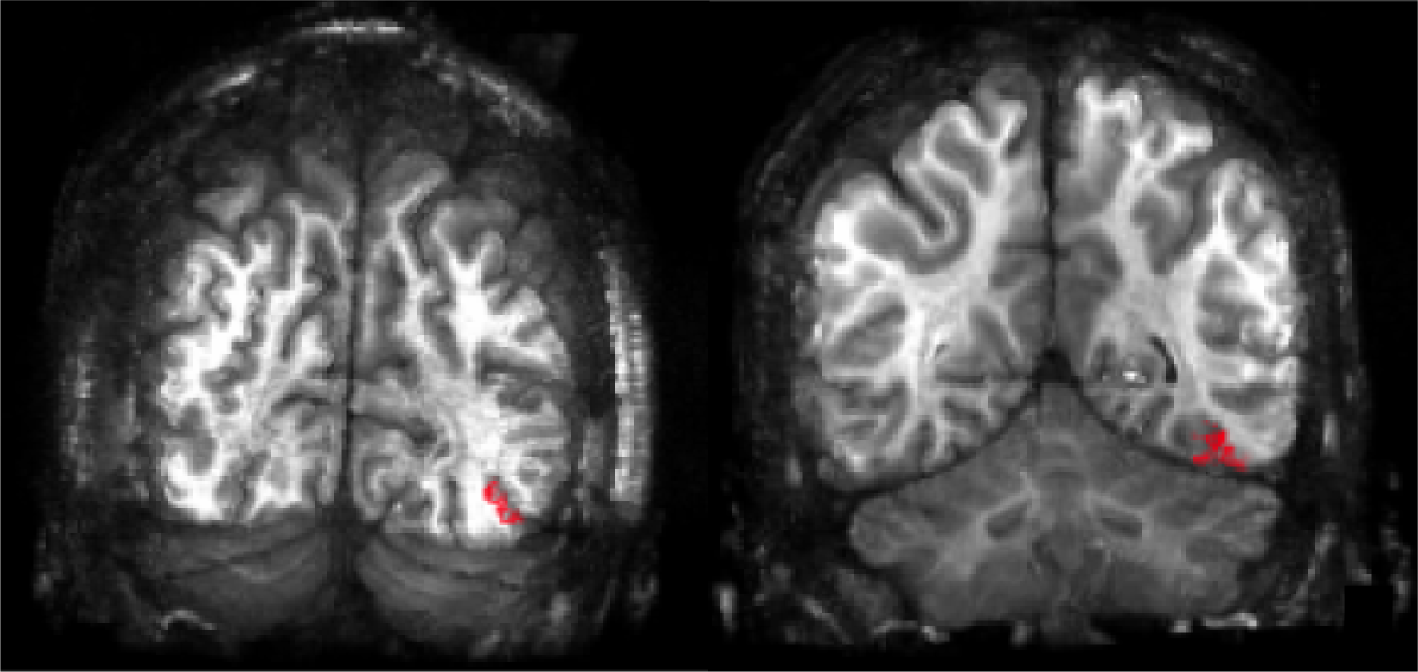
Functional ROI as defined in two subjects on IR-EPI.

### Depth dependent signal extraction

The fROI produced through the procedure described above was used to mask the layer-volume distribution. This resulted in a layer-volume distribution specific to the fROI. By treating this distribution as a design matrix such that rows were voxels and columns were bins, it was possible to regress it against the signal observed in the fROI for each time point in the experiment (van Mourik et al., 2019; Polimeni et al., 2010b).

Fitting the voxel-volume distribution to each time point in the experiment yielded the relative contribution of each bin to the overall signal at each time point, thereby representing a depth dependent time-series for each depth bin. These were treated similarly to voxel time-courses and used to fit the task design.

The task design model was then fitted to the extracted depth dependent time courses. Percent signal change was calculated as a division between *β*-weights assigned to each condition and the average weight assigned to the constant terms. The percent signal change values were then analyzed at the group level in an ANOVA and subsequent two-tailed, paired *t*-test comparing the responses to words and pseudo-words. *T*-statistics and ANOVA results were determined to be significant at *p* = 0.01. These analyses were implemented in MATLAB.

### Statistical analysis

#### Intraregional gPPI

The gPPI analysis is a generalized version of the PPI analysis. In the generalized form, the analysis is designed to span the full experiment space (McLaren et al., 2012). In gPPI analysis, the first level model is extended by including the time-course of a seed region in addition to interaction terms of the seed region with each task condition regressor.

As the goal of this analysis is to observe the effect of the interaction between the task and the neuronal response of a seed region, a deconvolution is typically applied to the seed time-course before computing the interaction term. Given the problems associated with deconvolution (O’Reilly et al., 2012) and the novel nature of this work, we omitted deconvolution from our gPPI analysis. The depth dependent hemodynamic response function (HRF) is not well understood. In the absence of this knowledge, the task regressors used in this experiment were created using the canonical HRF. Deconvolution based on the canonical HRF would have therefore exacerbated errors in modeling introduced by the initial convolution. Furthermore, the design of this study sequentially presented five items of each condition type, essentially in 20 second blocks. O’Reilly et al. (2012) have shown that omitting deconvolution is not expected to greatly affect the outcome in block designs such as that used in the present study. We therefore considered it more prudent to omit rather than include deconvolution in this analysis. There is no known precedent for laminar specific gPPI. The gPPI design was created by adding seed region time-courses and interaction terms to the original design. Interaction terms were calculated as the product of the detrended depth dependent time-series and binary condition vectors (1 when a condition response was expected, 0 when it was not) derived from the task regressors. A time point was included in the interaction term if the task regressor diverged from 0 by 0.0001.

Different models were created for each inter-regional analysis to assess the interaction between two depth-bins while alternating seed/target assignment. We did not model the third remaining bin. Six models were created in total, each containing six interaction terms (each of the six conditions multiplied by the seed-region time-course), the seed-region itself, and the full design as discussed previousy. Group effects were assessed using AFNI’s *3dANOVA3*. Paired two-tailed *t*-statistics were computed on the word and pseudo-word condition contrast. Results were deemed significant at *p* = 0.01.

#### Bin to whole brain gPPI

In a separate gPPI analysis, we modeled the task-dependent effect of the deep and middle bins on the whole brain. The superficial bin was excluded from the whole brain gPPI for several reasons. The IOTS is hypothesized to connect to left temporal cortex through either primarily bottom-up or top-down configurations (Price, 2012). To distinguish among these and thus explore the ability of lfMRI to distinguish between top-down from bottom-up network arrangements, it was necessary only to include the predicted top-down and bottom-up bins associated with word reading. Voxels within the superficial bin are also susceptible to partial volume artifacts due to vessels on the pial surface, which could possibly affect the analysis. The inclusion of the superficial bin would have required the gPPI model to include ten additional regression terms which would have been collinear with the experimentally interesting deep and middle bin terms, and so this was not considered further in the interests of a parsimonious data analysis.

The analysis was performed on the MNI normalized data with a 3mm Gaussian smoothing kernel applied. Alignment quality was assessed partially on the basis of alignment accuracy of the middle temporal gyrus. Individual subject *t*-statistic maps were used in conjunction with subject anatomy and the MNI template used for normalization to determine the alignment quality of task critical regions. Inaccuracies in subject registrations were addressed with a manually created, secondary transformation containing small translations intended to improve task critical region alignment without introducing large global inaccuracies. As the experimental question related to the depth dependent connectivity to regions that respond to the word/pseudo-word contrast, the use of the non-laminar first level maps to facilitate alignment was independent of the gPPI analysis. Group results were assessed using AFNI’s *3dANOVA3*, as in the previous section.

The parameters of the spatial AutoCorrelation Function (ACF) representing the smoothness of the data were computed using AFNI’s *3dFWHMx* on the residual time-series of first level analysis. The ACF parameters were used by the AFNI program *3dClustSim* to compute the likelihood of random clusters given the ACF parameters 0.4747, 3.2569, and 7.8772, in a volume of the dimensions 63 × 82 × 55 with 2mm isotropic voxels. The dimensions of the volume used for permutation testing were determined with a group level functional data mask. Clusters were deemed significant at *p*_uncorr_ = 0.001, *α* = 0.05.

The task-independent connectivity from the deep and middle bins was assessed using the deep and middle bin time-courses included in the gPPI model. The final results were visualized using the rendering plugin in AFNI. In addition to subjects 2 and 19, subject 4 was excluded from this analysis. We were unable to successfully bring subject 4 into MNI space. Large inaccuracies in the registration resulted in the exclusion of this subject from the whole brain analysis. This subject was included in all analyses performed in native space.

## Acknowledgments

The authors thank those who contributed to this work, especially research assistants and technical staff.

## Funding

Support was provided by the NWO Language in Interaction Gravitation grant.

## Author contributions

DS was the principal author and responsible for data acquisition, investigation, project administration, data curation, and formal analysis. TvM contributed methods and software essential to this effort and was consulted for data visualization. LJB contributed to the selection of scan protocol parameters, assisted with data reconstruction, and provided invaluable logistical support. KS was part of conceptualization and supervision for the early portion of this project. KW was responsible for supervision, conceptualization, project administration and contributed to the review and editing process of this manuscript. PH and DGN conceptualized and secured funding for this project, provided supervision and oversight, and contributed extensively to the editing process.

## Competing interests

The authors have no competing interests.

## Data and materials availability

Data will be made available upon reasonable request.

