## Supplemental Data 2 for "Laminar Specific fMRI Reveals Directed Interactions in Distributed Networks During Language Processing"

| | False font Short | False font Long |
|---|---|---|
| 1 | [illegible] | [illegible] |
| 2 | [illegible] | [illegible] |
| 3 | [illegible] | [illegible] |
| 4 | [illegible] | [illegible] |
| 5 | [illegible] | [illegible] |
| 6 | [illegible] | [illegible] |
| 7 | [illegible] | [illegible] |
| 8 | [illegible] | [illegible] |
| 9 | [illegible] | [illegible] |
| 10 | [illegible] | [illegible] |
| 11 | [illegible] | [illegible] |
| 12 | [illegible] | [illegible] |
| 13 | [illegible] | [illegible] |
| 14 | [illegible] | [illegible] |
| 15 | [illegible] | [illegible] |
| 16 | [illegible] | [illegible] |
| 17 | [illegible] | [illegible] |
| 18 | [illegible] | [illegible] |
| 19 | [illegible] | [illegible] |
| 20 | [illegible] | [illegible] |
| 21 | [illegible] | [illegible] |
| 22 | [illegible] | [illegible] |
| 23 | [illegible] | [illegible] |
| 24 | [illegible] | [illegible] |
| 25 | [illegible] | [illegible] |
| 26 | [illegible] | [illegible] |
| 27 | [illegible] | [illegible] |
| 28 | [illegible] | [illegible] |
| 29 | [illegible] | [illegible] |
| 30 | [illegible] | [illegible] |
| 31 | [illegible] | [illegible] |
| 32 | [illegible] | [illegible] |
| 33 | [illegible] | [illegible] |
| 34 | [illegible] | [illegible] |
| 35 | [illegible] | [illegible] |
| 36 | [illegible] | [illegible] |
| 37 | [illegible] | [illegible] |
| 38 | [illegible] | [illegible] |

| | | |
|---|---|---|
| 39 | [illegible] | [illegible] |
| 40 | [illegible] | [illegible] |
| 41 | [illegible] | [illegible] |
| 42 | [illegible] | [illegible] |
| 43 | [illegible] | [illegible] |
| 44 | [illegible] | [illegible] |
| 45 | [illegible] | [illegible] |
| 46 | [illegible] | [illegible] |
| 47 | [illegible] | [illegible] |
| 48 | [illegible] | [illegible] |
| 49 | [illegible] | [illegible] |
| 50 | [illegible] | [illegible] |
| 51 | [illegible] | [illegible] |
| 52 | [illegible] | [illegible] |
| 53 | [illegible] | [illegible] |
| 54 | [illegible] | [illegible] |
| 55 | [illegible] | [illegible] |
| 56 | [illegible] | [illegible] |
| 57 | [illegible] | [illegible] |
| 58 | [illegible] | [illegible] |
| 59 | [illegible] | [illegible] |
| 60 | [illegible] | [illegible] |
| 61 | [illegible] | [illegible] |
| 62 | [illegible] | [illegible] |
| 63 | [illegible] | [illegible] |
| 64 | [illegible] | [illegible] |
| 65 | [illegible] | [illegible] |
| 66 | [illegible] | [illegible] |
| 67 | [illegible] | [illegible] |
| 68 | [illegible] | [illegible] |
| 69 | [illegible] | [illegible] |
| 70 | [illegible] | [illegible] |
| 71 | [illegible] | [illegible] |
| 72 | [illegible] | [illegible] |
| 73 | [illegible] | [illegible] |
| 74 | [illegible] | [illegible] |
| 75 | [illegible] | [illegible] |
| 76 | [illegible] | [illegible] |
| 77 | [illegible] | [illegible] |
| 78 | [illegible] | [illegible] |

| | | |
|---|---|---|
| 79 | [illegible] | [illegible] |
| 80 | [illegible] | [illegible] |
| 81 | [illegible] | [illegible] |
| 82 | [illegible] | [illegible] |
| 83 | [illegible] | [illegible] |
| 84 | [illegible] | [illegible] |
| 85 | [illegible] | [illegible] |
| 86 | [illegible] | [illegible] |
| 87 | [illegible] | [illegible] |
| 88 | [illegible] | [illegible] |
| 89 | [illegible] | [illegible] |
| 90 | [illegible] | [illegible] |
| 91 | [illegible] | [illegible] |
| 92 | [illegible] | [illegible] |
| 93 | [illegible] | [illegible] |
| 94 | [illegible] | [illegible] |
| 95 | [illegible] | [illegible] |
| 96 | [illegible] | [illegible] |
| 97 | [illegible] | [illegible] |
| 98 | [illegible] | [illegible] |
| 99 | [illegible] | [illegible] |
| 100 | [illegible] | [illegible] |
| 101 | [illegible] | [illegible] |
| 102 | [illegible] | [illegible] |
| 103 | [illegible] | [illegible] |
| 104 | [illegible] | [illegible] |
| 105 | [illegible] | [illegible] |
| 106 | [illegible] | [illegible] |
| 107 | [illegible] | [illegible] |
| 108 | [illegible] | [illegible] |
| 109 | [illegible] | [illegible] |
| 110 | [illegible] | [illegible] |
| 111 | [illegible] | [illegible] |
| 112 | [illegible] | [illegible] |
| 113 | [illegible] | [illegible] |
| 114 | [illegible] | [illegible] |
| 115 | [illegible] | [illegible] |
| 116 | [illegible] | [illegible] |
| 117 | [illegible] | [illegible] |
| 118 | [illegible] | [illegible] |

119 ᑫᗯᓄᓯᒧ ᑫᚽᖆᓯᑕᐱᚽᚽᒣ

120 ᑎᐱᐱᓯ ᑎᚽᖆᑌᚽᖆᓯᑎᑕᐱᑌ
